# Gene loss associated with plasticity-first evolution in Heliconius butterflies

**DOI:** 10.1101/2025.06.22.660914

**Authors:** Erika C. Pinheiro de Castro, Francesco Cicconardi, Ian A. Warren, Nicol Rueda-M, Camilo Salazar, Søren Bak, Stephen H. Montgomery, Chris D. Jiggins

## Abstract

Phenotypic plasticity occurs when a genotype can produce more than one phenotype under different environmental conditions. Genetic accommodation allows plastic phenotypes to be tuned to new environments and eventually to even be lost. The colourful and toxic Heliconiini butterflies have biochemical plasticity – they either sequester their cyanogenic glucosides (CG) from their larval hostplant or biosynthesize them when compounds for sequestration are not available. Here, we trace the evolution of CG biosynthesis in Heliconiini butterflies, a fundamental component of this biochemical plasticity. We first reconstructed the evolutionary history of biochemical plasticity in Heliconiini butterflies using chemical data from over 700 individuals, demonstrating that plasticity was ancestral in the tribe and further lost in a few clades, such as the Sapho clade that only sequester CGs. In lepidopterans, CG biosynthesis has been fully characterized in the moth *Zygaena filipendulae*, and is composed of CYP405A2, CYP332A3, and UGT33A1. Thus, we CRISPR-edited the gene *CYP405* in *Heliconius erato,* and confirmed that *CYP405*-knockout caterpillars do not biosynthesize CGs. We identified the *CYP405*s and *CYP332*s in other lepidopterans and found that both genes were independently co-opted into CG biosynthesis in the Heliconiinae butterflies and *Zygaena* moths. Most lepidopterans have a *CYP332*, but *CYP405* is mostly restricted to butterflies and has been duplicated in all *Heliconius* species. Although several *CYP405* copies were found in the Sapho clade, they lack structurally important P450 domains, which explain they loss of biochemical plasticity via specialization in CG sequestration. Our findings represent one of the few examples of plasticity-first evolution in which the genetic mechanisms associated with its accommodation/assimilation are known.

## INTRODUCTION

Adaptative phenotypic plasticity allows species to maximise fitness across different environments, as a single genotype can produces multiple condition-dependent phenotypes. Such plasticity can also promote diversification under the ‘plasticity first’ hypothesis, in which ancestral plasticity facilitates the colonization of novel habitats and niches, but can be further lost as lineages specialize locally [1]. Although most fitness traits have some degree of plasticity, phenotypic plasticity has been mainly studied in morphological and physiological traits[2]. Chemical traits are fine-tuned by the environment and offer an excellent opportunity to exploit the molecular bases of phenotypic plasticity, promoting and maintaining diversity.

Chemical defences create an “enemy-free” niche for protected organisms to radiate and speciate [3]. Yet, chemically protected species need to evolve not only mechanisms to acquire toxins, but to also tolerate/resist them and both mechanisms need to be present for toxicity to translate into an adaptive advantage [4]. To make the evolutionary trajectory of chemical defences even more complicated, most mechanisms to acquire and tolerate toxins are biochemical pathways – such that all substrates, enzymes and intermediates need to be present for the toxin to be formed. Chemical defence is therefore a complex trait which may be difficult to evolve. As such, gene co-option [5] and horizontal gene-transfer [6,7] can facilitate the acquisition of toxicity. Furthermore, the evolution of toxicity and its complexity goes beyond genetics: For most heterotrophic organisms, diet also plays a crucial role, by either providing toxins that can be directly sequestered or the precursors for their biosynthesis [8]. This implies that: (i) the evolution of toxicity often cannot be separated from the evolution of diet specialization; and (ii) there is substantial phenotypic plasticity in animal toxicity in response to diet, although this is rarely considered as an example of adaptive plasticity.

Cyanide is a toxin that disrupts the mitochondrial electron transport chain and is a breakdown product of cyanogenic glucosides (CG). Some Lepidoptera have evolved to detoxify cyanide and feed on plants protected by CG[9]. While lepidopteran adaptations to detoxify cyanide came from both horizontal gene transfer (i.e. β-cyanoalanine synthase - [10] and co-option of genes with existing functions (i. e. Rhodanase - [11]), the evolution of the mechanisms used to acquire CG are less well understood. Possibly as a counter-adaptation against these specialized lepidopterans, some plant lineages/populations have disabled sequestration by changing the structure of their CGs - e. g. certain *Passiflora* species against heliconiine butterflies [12] - or even by becoming acyanogenic - e.g. *Turnera ulmifolia* in Mexico against *Euptoieta* butterflies [13]; and *Lotus corniculatus* against *Zygaena* moths [14]. These changes in the CG composition of the hostplant affect the biochemical phenotype of the lepidopteran that sequester these toxins. In return, some lepidopterans have evolved mechanisms to maintain their toxicity regardless of the CG profile of their host plants by biosynthesizing their own CGs.

Both higher plants and lepidopterans *de novo* biosynthesize CGs using the same three enzymatic steps: they convert amino acids into oximes and then into cyanohydrins, which are then stabilized with a glycosylation - forming a CG [15] (Figure 1). To assemble this pathway, they have recruited cytochromes P450 (P450s/CYPs) and UDP-glycosyltransferases (UGTs) (Figure 1). Despite having the same precursors, intermediates, products, and catalytic reactions, *de novo* biosynthesis of CGs evolved independently in lepidopterans and plants [15]. Moths of the family Zygaenidae and butterflies of the subfamily Heliconiinae are the only insects currently known to *de novo* biosynthesize CGs and both clades produce the same aliphatic compounds. In lepidopterans, CG biosynthesis has been fully characterized in the six-spot burnet moth *Zygaena filipendulae* [15], and is composed of CYP405A2, CYP332A3, and UGT33A1. Three *CYP405* and one *CYP332* homologues have been found in the genome of the butterfly *Heliconius melpomene*: *HmCYP405A5*, *HmCYP405A5, HmCYP405A6,* and *HmCYP332A1* [16]. These genes have all the conserved domains/motifs of a functional P450 and are expressed in life stages of intense CG biosynthesis [12,17], but their involvement in CG biosynthesis in heliconiines has not yet been empirically confirmed. Since moths and butterflies split ∼100 MYA [18], if CG biosynthesis in these two clades involves orthologous genes it is likely that these were independently co-opted to this pathway, but this hypothesis remains to be tested.

**Figure 1.**
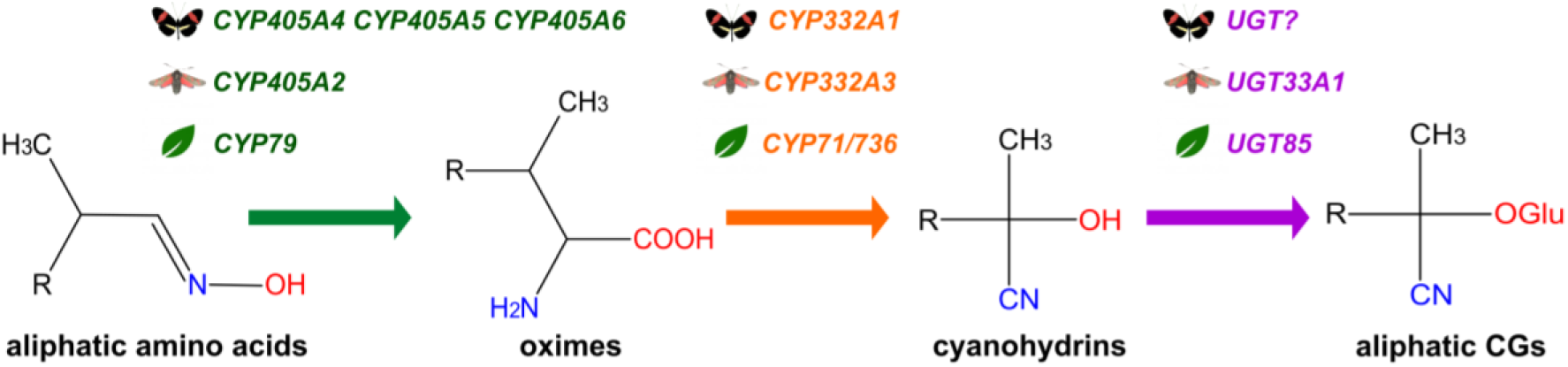
*De novo* biosynthesis of aliphatic cyanogenic glucosides (CGs) in *Heliconius melpomene* butterflies [16], *Zygaena filipendulae* moths [15], and plants, such as *Lotus* and cassava. The pathway in Heliconius was putative and enzymes were identified by homology with *Z. filipendulae*. The colours of the arrows correspond to the tree catalytic steps of CG biosynthesis and match the colour of the enzyme names that catalyse them. This pathway has the same enzymatic steps in both plants and lepidopterans because of convergent evolution [15]. The radical R in the chemical structures correspond to either CH_3_ or CH_2_-CH_3_.

While biosynthesis of CG allows them to maintain toxicity regardless of hostplant chemistry, it may have higher fitness costs than CG sequestration. Cyanogenic lepidopterans only biosynthesize CGs when they cannot sequester these compounds during larval feeding [19,20]. The CGs sequestered by heliconiines (cyclopentenic) have a different structure to those that are biosynthesized (aliphatic) [21]. The difference in structure between sequestered and synthesised compounds enables the use of target metabolomics to identify which strategy wild-caught heliconiine individuals used to acquire their CG reservoir [22]. This creates a unique opportunity to study the importance of phenotypic plasticity in the evolution of toxicity. Moreover, phenotypic plasticity can itself evolve when selection acts to change reaction norms. The different degrees of hostplant specialization within Heliconiini, in which different clades specialize on specific subgenera of *Passiflora*, with some feeding on several hostplants, while others are completely monophagous, creates a unique system in which to understand how specialization affects biochemical plasticity [23]. Finally, heliconiine butterflies have been studied for over 150 years, with recent efforts resulting in assembled reference genomes [24] and target-metabolomics data for most species of the tribe [22,25], which allows for powerful comparative analyses to reveal the genetic basis of the maintenance and loss of biochemical plasticity.

We first used target-metabolomic data for over 700 individuals [22,25] to evaluate the importance of biosynthesis versus sequestration in each clade and reconstruct the ancestral path of CG acquisition. We then used the gene-editing technology CRISPR-Cas9 to confirm the role of *CYP405As* in CG biosynthesis in *Heliconius*. Next, we explored the evolutionary history of these genes across 58 heliconiine genomes [24] and used molecular evolution analysis to test for selection on specific branches of the phylogenetic tree. We hypothesize that *CYP405s* are only duplicated in *Heliconius* which are more toxic than other heliconiines and that these copies might have been lost in some *Heliconius* clades that have specialized in sequestration and have lost biochemical plasticity.

## METHODS

### CRISPR-editing of *CYP405As* in *Heliconius erato demophoon*

CRISPR-Cas9 gene editing was used to confirm that *CYP405A*, previously characterized in Z*ygaena* moths, is also involved in the *de novo* biosynthesis of CG in heliconiines. This experiment was performed using a population of *H. erato demophoon* kept under insectary conditions (Supplementary Methods) This species has three copies of *CYP405As*: *HedCYP40A5A4*, *HedCYP40A5A6a* and *HedCYP405A6b*. Single-guides RNAs (sgRNA) targeting these genes were designed using the software Geneious (v2020.1) by identifying N20NGG sites within exons of the target-genes and internally blasting these sequences against the genome of *H. erato demophoon* v1 (http://ensembl.lepbase.org/Heliconius_erato_demophoon_v1/Info/Index) to remove sgRNA candidates with possible off-targeting sites. sgRNA4 ( GCCTAACGGTGATCCTCAGAAGG) was selected targeting a region in Exon 7 of *HedCYP40A5A4,* and sgRNA6ab ( GCCTAACGGTGACCCCATGAAGG) targeting the same region in both *HedCYP40A5A6a* and *HedCYP405A6b.* Unfortunately, it was not possible to design a sgRNA that could distinguish between the CDS of these two orthologues due to their similarity. sgRNAs were ordered from Synthego (https://www.synthego.com/) and dissolved following the manufacturer’s instructions. Freshly-laid eggs of *H. erato* (within 1-2h of oviposition) were glued on microscope slides and injected with a mix containing 200 ng Cas9 (TrueCut Cas9 ProteinV2, Invitrogen) with 200 ng sgRNA per µL. As a control for each batch, some eggs were glued on microscope slides and injected with Cas9 only (200 ng/ul) or kept non-injected. Larvae from control and CRISPR-eggs were individually raised in tubes and fed *ad libitium* with *Passiflora biflora* leaves, which does not contain CGs that can be sequestered by heliconiines (forcing them to biosynthesize their own CGs). Due to high mortality in the experimental group (injected with the sgRNAs), larvae were sampled after 7-8d of hatching. For the phenotypic analyses, larvae were weighed at the day of sampling and part of their tissue used for quantification of CGs using LC-MS following the extraction and analytical methods described in [12]. ANOVA was performed in R (R Core Team, 2018) to test if the weight and CG content of CRISPR-edited larvae was lower than the control.

### Genotyping the CRISPR-edited CYP405s and control larvae

Nanopore Minion sequencing of PCR products was used for genotyping using the four primer PCR kit (Oxford Nanopore Technologies plc., Oxford UK, Kit: SQK-PBK004). DNA was extracted using an in house bead extraction protocol ([26]). The following primers were used to amplify DNA from all three copies of the gene and add adapters for sequencing (Forward: TTTCTGTTGGTGCTGATATTGCAAGATTGACTCCATTGCACACT, Reverse : ACTTGCCTGTCGCTCTATCTTCCAATGACATCAGGTACATTGCG). PCR products were visualised on an agarose gel prior to sequencing library preparation and barcoding, which was performed using the manufacturer’s protocol (kit: SQK-PBK004). Sequencing was carried out until each library had 10,000 reads. Basecalling was done by the guppy basecaller (Wick et al., 2019), adapter trimming and quality control was performed using pore-chopper (https://github.com/rrwick/Porechop) and quality control was performed using MinionQC.r [28]. Minimap2 (Li, 2018) was used to align the reads to a contig containing all three gene copies. Samtools [30] was used to convert resulting output to a BAM file, and then to sort and index the files. The alignments were visualized in Geneious v2020.1.

### Biosynthesis versus sequestration profile of heliconiines and ancestral reconstruction analyses

Published datasets from [12], [31] and [32] on the CG profile of different heliconiine species were used for this analysis, comprising 743 butterflies of 43 species. The concentration of each CG (µg/mg), as well as the total CG concentration (µg/mg), found in each examined individual was extracted from the original datasets. The aliphatic CGs linamarin, lotaustralin and epilostaustralin were labelled as ‘biosynthesized CG’ (Table S1). The cyclopentenyl CGs deidaclin/tetraphyllin A, tetraphyllin B, epivolkenin, gynocardin, and dyhidrogynocardin (Table S1) were labelled as ‘sequestered CGs’ (Table S1). The sum of the concentration of all biosynthesized CGs and all sequestered CGs in each individual was divided by its total CG concentration to calculate ‘% of biosynthesis and % of sequestration, respectively. An average of % of biosynthesis and % of sequestration for each species was calculated using all related individual entries and plotted in bi-directional bar charts per species following the tree hypothesis from [24]. This dataset was also used to perform ancestral reconstruction analysis on biochemical plasticity, using the function “anc.ML” the package “Phytools”[33].

### Cytochrome P450 CYP405s and CYP332: phylogeny, convergence evolution & evolutionary pressure analyses

To evaluate the phylogenetic relationship of *CYP405s* and *CYP332s* and test whether the co-opted genes in the moth *Zygaena filipendulae* and the butterfly subfamily Heliconiinae belonged to the same orthogroups (OGs), we first downloaded all P450 genes of the CYP4 and CYP3 clades from insectp450.net database [34] and selected all sequences from Lepidoptera annotated as complete, for a total of 9230 sequences from 26 families. Amino-acid sequences were aligned with ClustalW and used to generate the full maximum likelihood phylogenetic tree using IqTree2 [35] using the LG+G4+F and 1000 ultrafast bootstrap replicates. Once the OGs were identified, we added all those genes to our list to generate the full OG phylogenetic trees using nucleotide sequences (see below). The protein sequence of CYP405s and CYP332s from *Heliconius melpomene* and *H. erato* annotations were obtained from NCBI, as well as the sequences from *Bombyx mori*. All these sequences were used as queries in EXONERATE[36] [setting: -n 3 --model protein2genome --percent 50 --refine full] to annotate coding exons and introns in 58 heliconiini and other two heliconiinae (*Speyeria mormonia* and *Brenthis ino*) species. The resultant nucleotide sequences were subsequently mapped using MINIMAP2(LI, 2018) to explore genome assemblies where no record was found in the EXONERATE mapping to minimise false negative hits. The results were used as a template to manually check and correct the sequences in all species. All the amino acid sequences obtained were processed with CDSEARCH webserver[38] and BLASTP webserver [39] to verify the presence and the completeness of the conserved domain (PFAM00067) and possible gene-fusions. All nucleotide sequences were aligned using MACSE2 [40], and an initial maximum likelihood phylogenetic gene tree reconstruction was generated as implemented in IQ-TREE2 [-M MFP -B 5000]. At this point all sequences with putative stop-codons were removed from the alignment and ML gene tree recalculated and used as input for follow-up analyses.

To evaluate convergent evolution in the two orthogroups in *Zygaena* moth and Heliconiinae butterflies, we used CSUBST [41] to derive observed non-synonymous convergence (OCN), observed synonymous convergence (OCS), and the relative ωC metric ((OCN/ECN)/(OCS/ECS)), being ECN and ECS, the expected non-synonymous and synonymous substitutions. We ran the analysis accepting all convergent events that occurred in all versus all lineage comparisons without specifying any lineages as foreground and background. From the analysis we evaluated ωC, OCN and OCS, each with a cutoff of 1. Differences in the distributions were tested using one-sided Wilcoxon rank-sum tests.

Several tests for selective forces were carried out for *CYP405* and *CYP332* in the tribe Heliconiini: Initially, all sequences from each locus were realigned separately, and model test and the overall *ω* (hyphy acd Universal $FASTA MG94CUSTOMCF3X4 Global) were calculated as implemented in HYPHY v2.5.32(MP) for Linux on x86_64 [42] batch language. In addition, the Single-Likelihood Ancestor Counting (SLAC) method [43], as implemented in the HYPHY batch language, was used to evaluate the total number of sites under purifying and positive selection, with a *P* value of 0.05 as threshold.

Previous studies on the CG profile of heliconiines, i.e. studies [12] and [31] found that some *Heliconius* species in the Sara-sapho clade have specialized on sequestration and might even have lost their biosynthetic ability (e.g. *H. sapho*, *H. antiochus* and *H hewitsoni*). In order to detect possible shifts in the selection acting upon these biosynthetic genes in this group, two tests were performed: First, we applied BUSTED-PHENOTYPE (BUSTED-PH.bf at https://github.com/veg/hyphy-analyses), a method that detects evidence of episodic diversifying selection associated with feature, phenotype or trait. This is built on BUSTED [47] and it is implemented under HYPHY batch language. BUSTED-PHENOTYPE can detect evidence of episodic diversifying selection within groups and between groups. Nevertheless, because BUSTED-PHENOTYPE does not describe the direction of the selection, we performed the RELAX [44] test to investigate whether the phenotype/trait branches experienced relaxation of intensification, and thus the direction of the selection.

## RESULTS

### Variation in the balance of CG biosynthesis and sequestration across the phylogeny

We first reconstructed the evolutionary history of biochemical plasticity in the tribe Heliconiini using a phylogenetic approach, using published datasets on the cyanogenic glucoside (CG) profile of 743 heliconiine butterflies (Figure 2). At the base of the heliconiine phylogeny (Figure 2, “Other genera”) CG biosynthesis is the dominant strategy. Nevertheless, sequestered CGs were found in a few individuals of all basal species (*Philaethria dido*: 1 out of 6*; Dryas iulia:* 15 of 45; *Agraulis vanillae:* 7 out 12; and *Dryadula phaetusa:* 1 out of 15), except for *Dione juno* (0 out of 15). This suggests that although biosynthesis is the dominant strategy, the ancestral state involved some degree of plasticity. Similarly, plasticity was also present in the ancestor of the genus *Eueides,* but biosynthesis is the most common strategy, with sequestration observed in only two *Eueides isabella* butterflies out of 34 *Eueides* individuals sampled, and in none of the other four species analysed.

**Figure 2.**
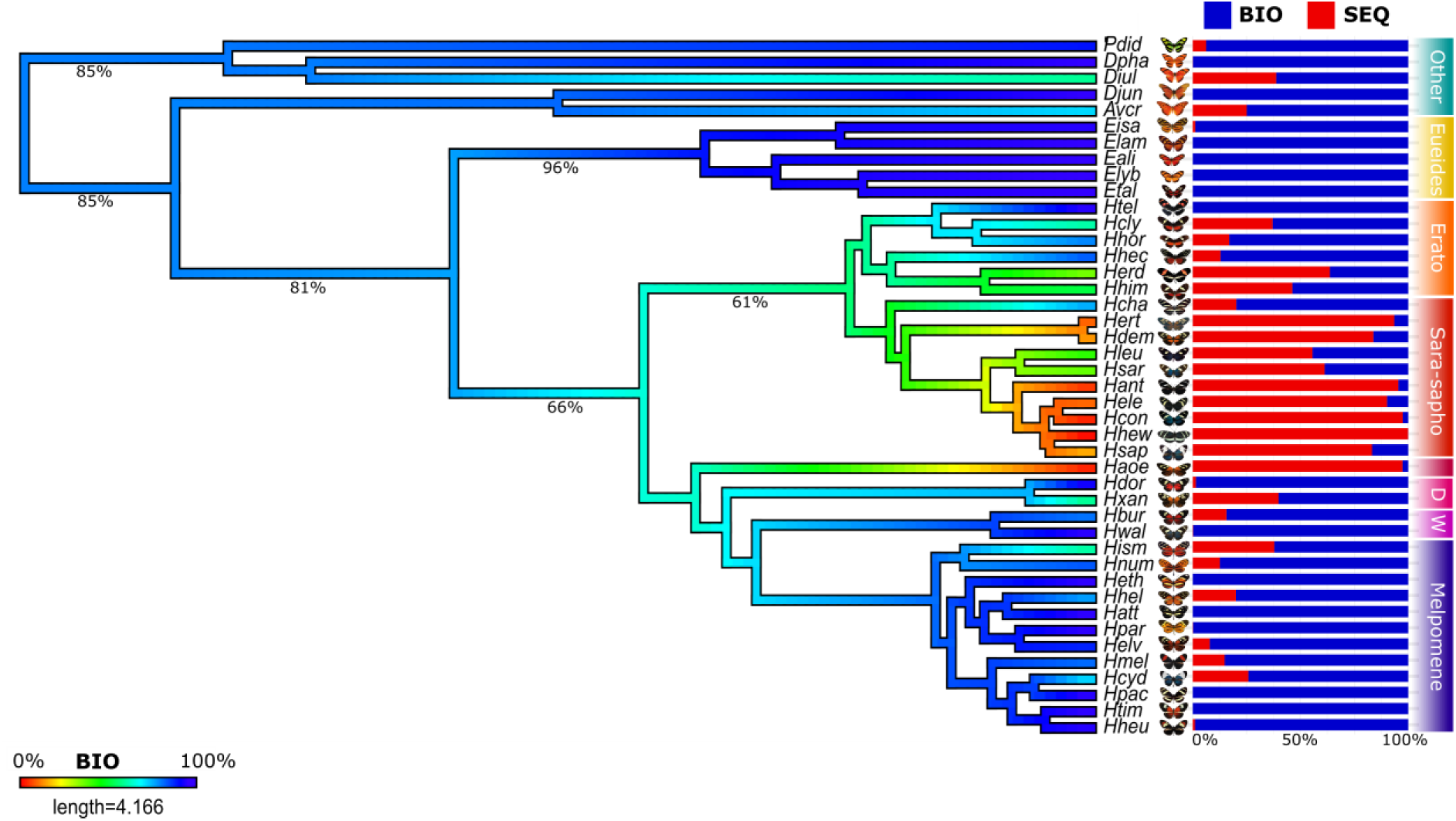
Ancestral reconstruction of biochemical plasticity in the Heliconiini tribe. This analysis used the tree hypothesis from Cicconard et al. 2023 and a chemical dataset (743 butterflies) built based on the raw data from Castro et al. 2018, Sculffold et al. 2020, and Rueda et al. 2023). Bars correspond to average percentage of biosynthesized (blue) versus sequestered (red) CGs per species and colour of the branches to estimated percentage of biosynthesized CGs (from 0% in red to 100% in blue) in the tree internal nodes. Species names are highlighted according to their clades. Legend: OTHER genera - *Pdid= Philaethria dido; Dpha= Dryadula phaetusa; Diul= Dryas iulia; Djun= Dione juno; Avcr: Agraulis vanillae. genus EUEIDES - Eisa= E.isabella; Elam: E. lampeto; Eali: E. aliphera; Elyb= E. lybia; Etal= E. tales. genus Heliconius: ERAT0 – Htel= H. telesiphe; Hcly= H. clysonymus; Hhor= H. hortense; Hhec= H. hecalesia; Herd= H. erato; Hhim= H. himera; SARA-SAPHO: Hcha= H. charithonia; Hert= H. eratosignis; Hdem= H. demeter; Hleu= H. leucadia; Hsar= H. sara; H.ant= H. antiochus; Hele= H. eleuchia; Hcon: H. congener; Hhew: H. hewitsoni; Hsap= H. sapho. AOEDE: Haoe: H. aoede; DORIS: Hdor= H. doris; Hxan= H. xanthocles; WALLACEI: Hbur= H. burney; H. wallacei; MELPOMENE: Hism= H. ismenius; Hnum= H. numata; Heth= H. ethila; Hhel= H. hecale; Hatt= H. atthis; Hpar= H. pardalinus; Helv= H. elevatus; Hmel=H. melpomene; Hcyd= H. cydno; Hpac= H. pachinus; Htim= H. timareta; Hheu= H. heurippa*.

CG sequestration is more frequent in the ancestor of *Heliconius,* the most speciose genus of the tribe. CG sequestration was observed in almost all species within the Erato + Sara-Sapho clades, except *H. telesiphe* (out of 16 species). Indeed, the ancestor of the Sara-sapho clade favoured CG sequestration over biosynthesis and in some species, biosynthesis is rarely seen (i.e. *H. sapho*, *H. hewitsoni*, and *H. antiochus*). CG sequestration was also common in the MRCA Aoede + Doris + Wallacei + Melpomene. *Heliconius aoede* mostly contains sequestered compounds (small amounts of biosynthesis was found in 4 individuals out of 25). Nevertheless, CG biosynthesis regains importance in the MRCA Doris + Wallacei + Melpomene, especially in the Melpomene clade, in which some species (4 out of 12 – i.e. *H. pachinus*, *H. timareta*, *H. ethila* and *H. atthis*) only biosynthesize CGs and appear to have lost biochemical plasticity. Sequestered compounds were also virtually absent in *H. doris* (N=35) and *H. wallacei* (n= 14). In summary, from a highly plastic ancestor, some species within the genus *Heliconius* have evolved to specialise on either sequestration or biosynthesis of their CGs.

### H. erato with CRISPR-edited CYP405s do not biosynthesize CGs

We next used gene knockout experiments with CRISPR/Cas9 to confirm the role of *CYP405* genes in CG biosynthesis. Fresh eggs of *H. erato demophoon* were injected with a sgRNA targeting either *HedCYP405A4* or the duplicated *HedCYP405A6* (experimental group), with controls injected with just the Cas9 enzyme. Individuals in the experimental group showed reduced weight as compared to the control in the first week (Kruskal-Wallis, *X*^2^ = 38.065, Df=2, p= 5.423e-09) and most failed to develop to final instar (Figure 3A). Target-metabolomics showed that larvae with a deletion in the CDS of *CYP405A4 or CYP405A6s* lacked or had very low biosynthesized CG in comparison with others (Figure 3B). Larvae in the experimental group that did not have deletions in the *CYP405A4 or CYP405A6s* also had more CG than those in which gene-editing was successful. Nanopore sequence of CRISPR-edited and control individuals also confirmed that the designed sgRNAs were specific and only caused deletions in the target *CYP405* orthologue (Figure S1)

**Figure 3.**
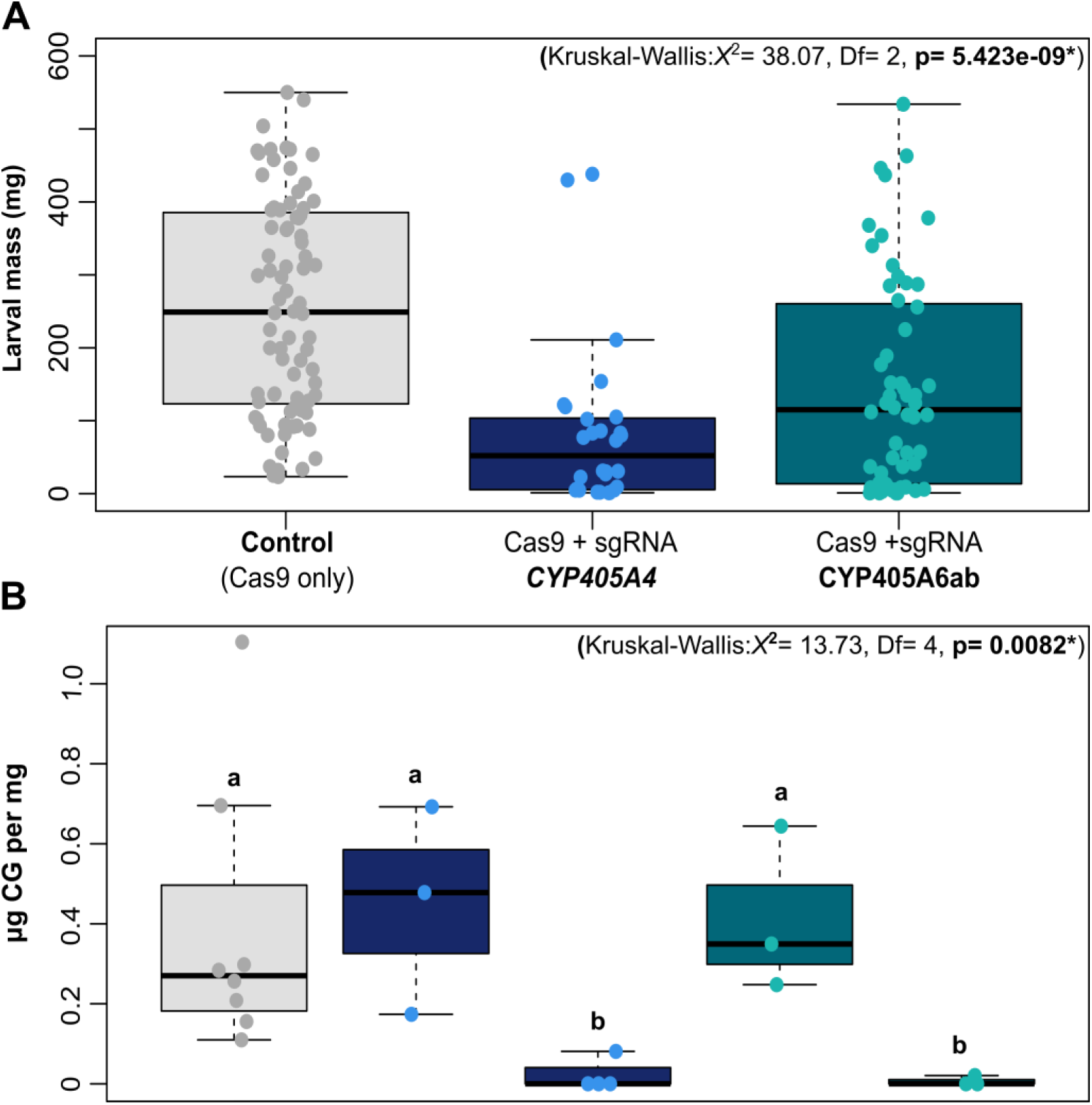
A) Larval mass (mg) of all 7d old larvae from injected eggs of *H. erato*. **B)** CG concentration (ug/mg) of genotyped larvae. Controls (grey) were injected with Cas9 only, whereas others were injected with Cas9 and the sgRNA specifically targeting either *CYP405A4* (navy blue) or *CYP405A6* (dark cyan). *CYP405A* knockouts had a deletion in the target copies, whereas not-edited larvae had the same genotype as control, even though they were injected with the respective sgRNAs. The specificity of the sgRNA used

### CYP405 and CYP332 were co-opted twice into CG biosynthesis in Lepidoptera

To evaluate whether *CYP405s*, as well as *CYP332s,* were independently co-opted into CG biosynthesis in butterflies and moths, we first used the available *Heliconius* CYP405A4 and CYP332A1 protein sequences as a starting query to identify homologues across 63 Heliconiinae genomes, identifying 144 *CYP405* and 66 *CYP332* loci (Table S2). Then, we constructed a maximum likelihood tree using these Heliconiinae sequences, as well as the amino acid sequences of all complete cytochromes P450 from clan CYP3 and CYP4 in lepidopterans (Figure S2). Whereas *CYP332* is present in most lepidopterans (Figure 5), *CYP405* is found only in a few moth species (Figure 4). All *CYP405s* and *CYP332s* in the subfamily Heliconiinae are very conserved and are monophyletic relative to other Lepidoptera. The two *CYP405s* from *Zygaena filipendulae* and CYP332 cluster as sister clades to butterflies instead of with other moths, although bootstrap support is relatively low. CSUBST analyses were performed to explicitly test for convergence in the protein sequences between *Zygaena* moths and Heliconinae butterflies and supported the hypothesis that both *CYP405* and *CYP332* where independently co-opted twice into CG biosynthesis in these two distinct lepidoptera lineages (Figure S3).

**Figure 4.**
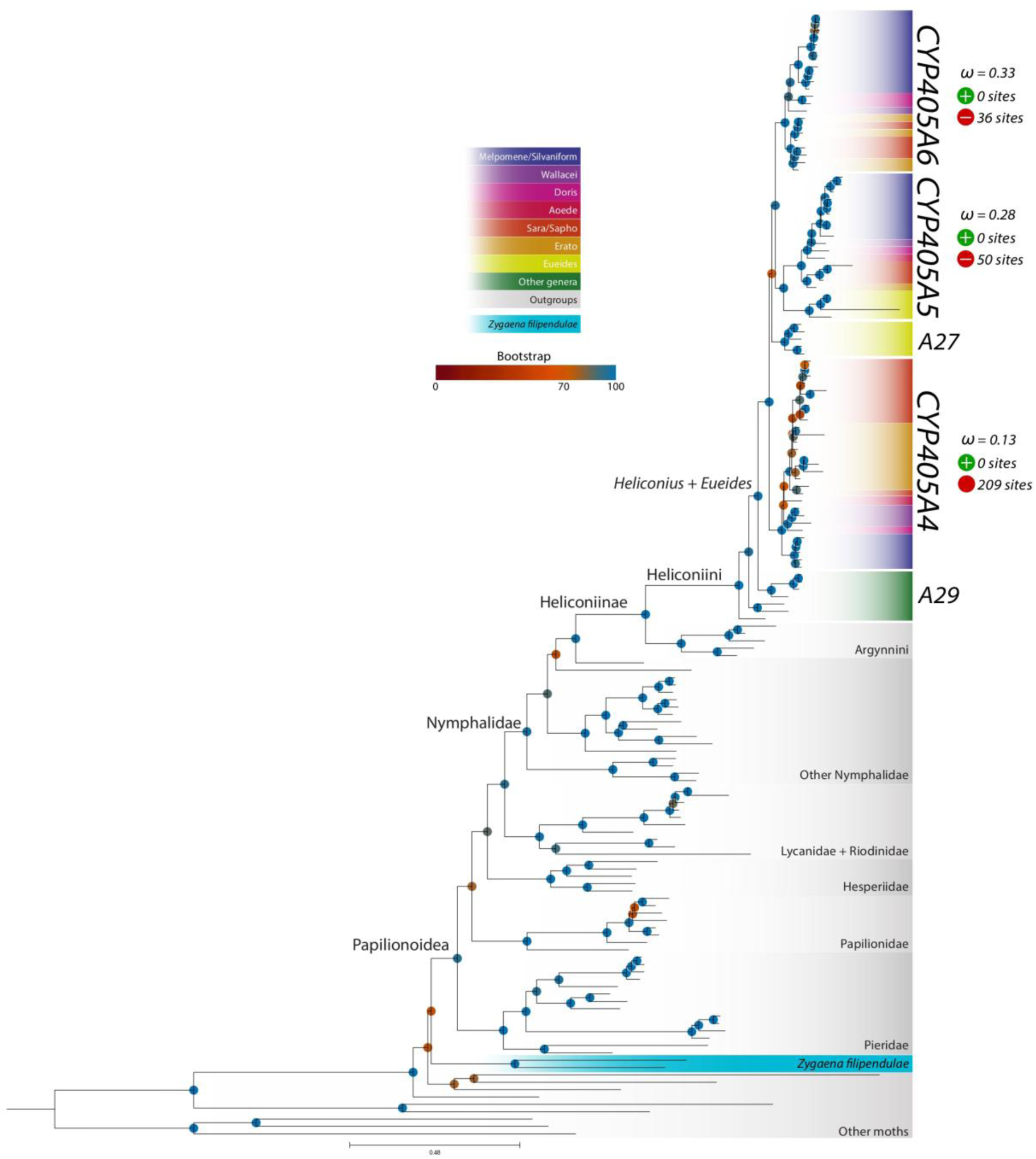
Gene-tree of CYP405A in Lepidoptera. Cyanogenic species (i.e. Heliconiini butterflies and *Zygaena* moths) are coloured per clade according to the figure legend and other lepidopterans in grey. Coloured circles correspond to branch bootstrap values (Red=low, Blue=high). Molecular evolution analyses were performed in the Heliconiini tribe, in which purifying selection (ω) on each CYP405 orthologues was quantified, as well as number of sites under purifying (negative) and diversifying (positive) selection identified.

**Figure 5.**
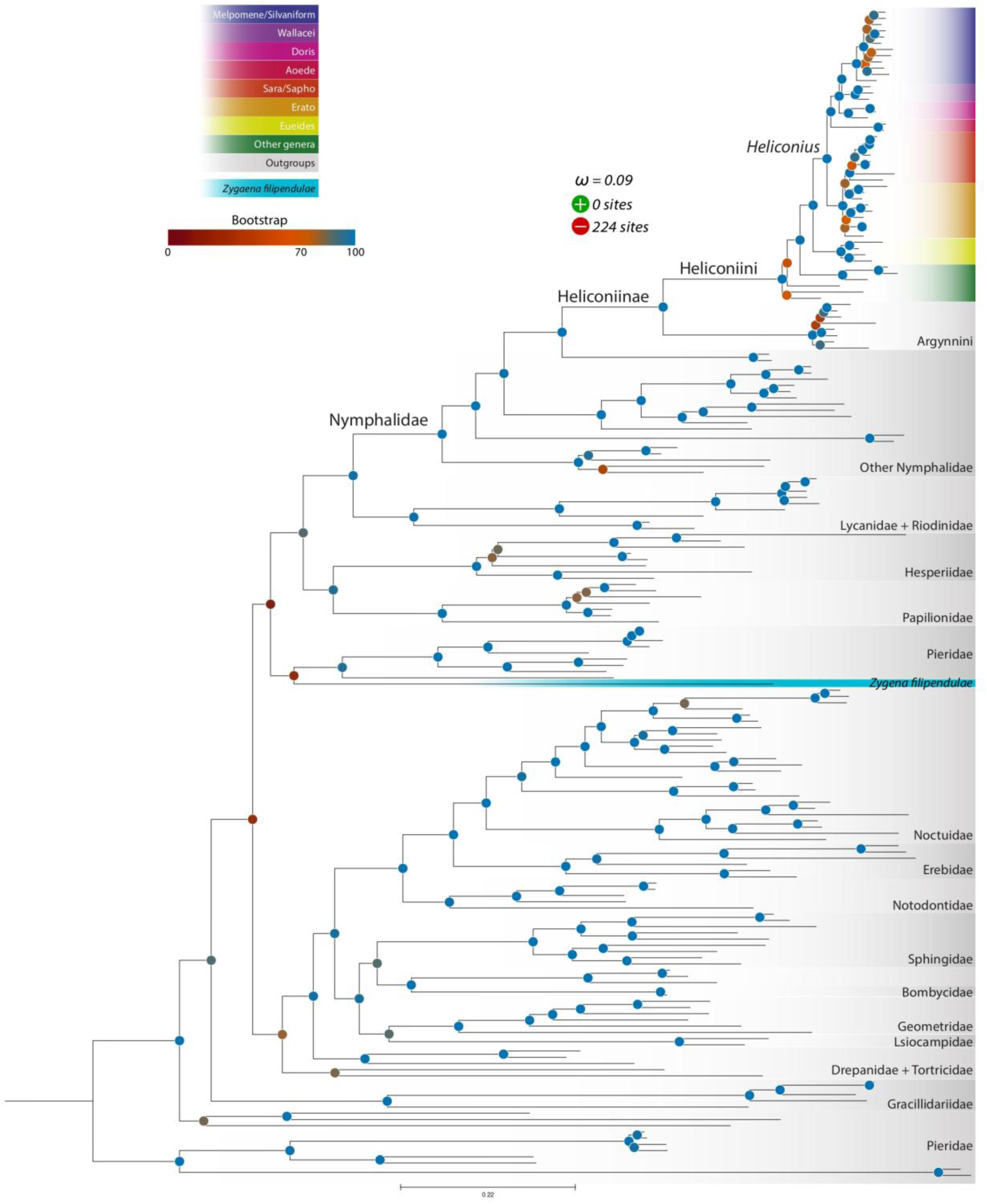
Gene-tree of CYP332 in Lepidoptera. Cyanogenic species (i.e. Heliconiini butterflies and *Zygaena* moths) are coloured per clade according to the figure legend and other lepidopterans in grey. Coloured circles correspond to branch bootstrap values (Red=low, Blue=high). Molecular evolution analyses were performed in the Heliconiini tribe, in which purifying selection (ω), as well as number of sites under purifying (negative) and diversifying (positive) selection identified.

### CYP405s duplicated in Heliconius and Eueides, but not in other heliconiines

*CYP405* was duplicated in the genera *Heliconius* and *Eueides*, but not in the other Heliconiini species (Figure 4). A few independent duplications and losses have occurred in specific *Heliconiu*s clades/taxa, but overall, most *Heliconius* species have three or more *CYP405*s. We hypothesize that an ancestral *CYP405* loci was present in the outgroups (*CYP405A30*), other Heliconiini (*CYP405A29*), *Dione/Agraulis* (*CYP405A28*) and *Heliconius* (*CYP405A4*) (Figure 4). The ancestral locus seems to have undergone a duplication (*CYP405A5*) at the MRCA *Heliconius* + *Eueides*, followed by a loss in all the species of the *Erato* clade and in half of the 10 *Sara-Sapho* clade species. The ancestral locus likely duplicated a second time at the stem of all *Heliconius* (*CYP405A6*). The synteny of *CYP405* is highly conserved with all loci located on the ancestral chromosome 15 (which corresponds to chromosome 17 in *Heliconius*).

*CYP332* is the second enzyme of the CG biosynthetic pathway, and two copies of this gene were found in *Speyeria mormonia* and *Brenthis* (Figure 5). The majority of Heliconiini have a single *CYP332* locus, with the exception of *H. hortense* and *H. hermethena* (within the Erato clade), *H. aoede*, and *H. elevatus*, which have two. The locus is located on the ancestral chromosome 3 (which correspond to chromosome 11 in *Heliconius*), without any evidence of translocations and/or inversions.

### Association between CYP405A-copy number, chemical phenotype and hostplant specialization

*Heliconius* species have up to four *CYP405* genes, but some sequences were smaller than expected (∼1500 bp). Translated sequences were therefore manually checked for the presence of the six conserved motifs in insect P450s (Feyereisen, 1999): Cluster of Pro/Gly, C-Helix, I-Helix, ExLR, PERF, and Heme-binding loop. This resulted in 98 complete *CYP405s* out of 144, with most of the incomplete copies found in the *Heliconius sapho clade* (14 incomplete sequences) (Figure 6).

**Figure 6.**
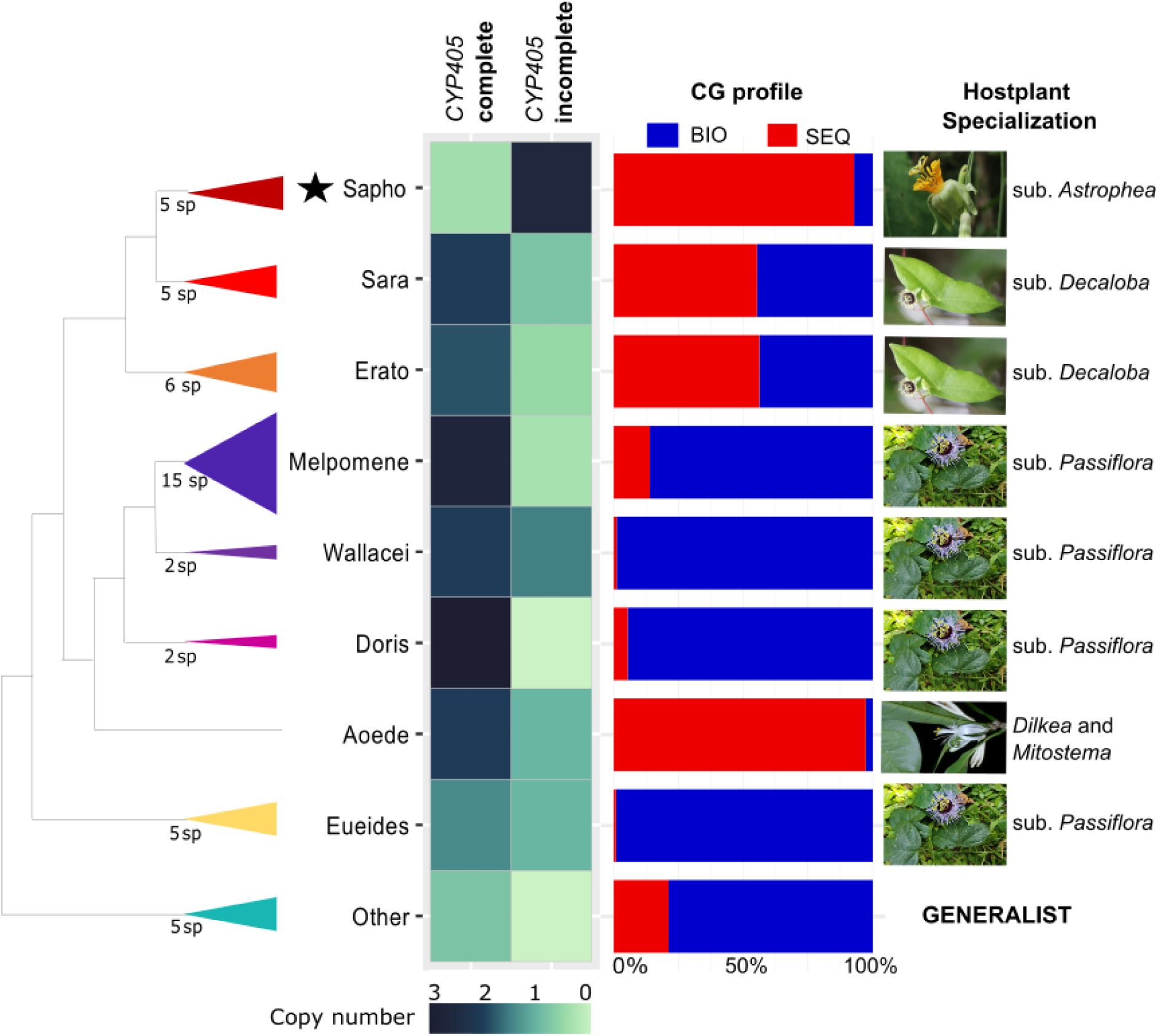
*CYP405*-copy number, chemical profile and hostplant specialisation of different Heliconiini clades. In this figure, the clade Sara-sapho was split into two distinct clades: Sara (*H. sara*, *H. leucadia*, *H. demeter*, *H. eratosignis* and *H. charithonia*) and Sapho (*H. sapho*, *H. antiochus, H. eleuchia, H. congener* and *H. hewitsoni*) , as they have different hostplant specialization, different chemistry and different number of complete *CYP405s*.

There is a clade-specific pattern in *CYP405*-copy number with the majority of the Silvaniform/Melpomene clade having three complete *CYP405s*, most Erato and Sara having two and Sapho none (Figure 6). This pattern matches the chemical profile of these clades. CG sequestration in the Heliconiini tribe gained more importance in the MRCA *Heliconius* Erato + Sara + Sapho, especially for Sapho species, and in the Aoede clade. This clade-specific pattern is also associated with their hostplant specialization: while other *Heliconius* (MRCA Silvanifom/Melpomene + Doris + Wallacei) have hostplants in the subgenus *Passiflora,* in which most species lack CGs that can be sequestered [22], the Erato and Sara clades have hostplants in the subgenus *Decaloba* and Sapho in the subgenus *Astrophea,* where CG for sequestration is common. Aoede species use the genera *Mitostemma* and *Dilkea* as hostplants from which they sequester CGs, instead of *Passiflora.* Our results therefore suggest that the loss of plasticity has also been associated with a loss of complete copies of the genes required for biosynthesis.

### P450s involved in CG biosynthesis are under purifying selection

Adaptative plasticity might be maintained by purifying section, therefore we quantified the levels of purifying selection acting across the *CYP405* orthologues and *CYP332.* We found that *ω* varies substantially among orthologues (Figure 4 and 5). The ancestral *CYP405A* locus, *CYP405A28-30* in the outgroups and other Heliconiini and *CYP405A4* in *Heliconius*, show purifying selection (*ω* = 0.13 and *ω* = 0.17). None of the orthologues had sites under positive selection. *CYP332* shows a stronger signature of purifying selection (*ω* = 0.09; 224 sites under purifying selection; no sites under positive selection) than the *CYP405As*.

We next tested whether “sequestration-specialist” species (part of the Sara-sapho clade and *Heliconius aoede*), show evidence for reduced purifying selection on genes involved in CG biosynthesis. BUSTED-PH was used to test whether *CYP405s and CYP332s* have experienced episodic positive diversifying selection in “sequestration-specialist” in comparison to the others (“plastic species”), and the RELEX test infers the direction of the shift between these two groups. For *CYP405,* we performed these analyses separately for each orthologue and found that in all orthologues except *CYP4056* episodic diversifying positive selection were detected in both “sequestration-specialist” and “plastic-species” , but no significant differences were detected. For *CYP405A6* episodic selection was not found in “sequestration-specialist”, but no significant difference was detected by BUSTED-PH in comparison with “plastic-species”. In the RELAX test, neither intensification nor relaxation were detected for each OG. For *CYP332*, only the “plastic-species” showed episodic diversifying positive selection, but no evidence of differential selection regimes nor intensification or relaxation of selection was found (P value < 0.0001) in the RELAX test .

## DISCUSSION

Phenotypic plasticity plays an important role in allowing organisms adapt to novel or extreme environments. However, plasticity is mainly studied in visible phenotypes. Our findings emphasize the importance of phenotypic plasticity for the evolution of toxicity, a complex molecular trait. Here, we first demonstrated, using a phylogenetic reconstruction, that there is a pattern of ancestral biochemical plasticity and derived loss of plasticity in Heliconiini butterflies, providing a novel example of plasticity-first evolution. We also use CRISPR to characterize *CYP405s* in *Heliconius erato*, demonstrating that these genes are involved in CG biosynthesis, which enables biochemical plasticity. Finally, we have shown that gene-loss is associated with the loss of plasticity in a CG sequestration-specialist clade (Sapho), demonstrating a genetic mechanism for plasticity-first evolution.

Passion vine butterflies and burnet-moths are the only lepidopterans known to biosynthesize CGs[9]. Yet, most butterflies and some moths have an extra copy of *CYP405*, suggesting that this gene was independently co-opted as the first-step of CG *de novo* biosynthesis at least twice and likely has another function in other Lepidoptera [49]. The high mortality and poor development of the *CYP405*-knockout larvae in our experiments might be associated with this other function and/or the importance of CG biosynthesis for nitrogen metabolism. Similar to lepidopterans, several distantly-related cyanogenic plants have co-opted *CYP79s* into the first step of CG biosynthesis [50], while ferns have recruited a FMO [51]. Indeed, although plants and lepidopterans biosynthesize CGs using the same enzymatic steps, the genes involved are not homologous and have evolved independently. In higher plants, different P450s have been recruited into the second step of CG biosynthesis [52], [53]. In lepidopterans, this second step is catalysed by CYP332, a gene characterized in *Zygaena* moths and found across all Heliconiini butterflies. While the ancestral function of *CYP405* in lepidopterans remains unknown, *CYP332* was likely involved in detoxification of plant toxins[49]. Most lepidopterans have one *CYP332,* but interestingly, *Helicoverpa armigera* has two copies: while *CYP332A1* has undergone neofunctionalization and is associated with resistance to fenvalerate, the function of *CYP332A2* is still unknown [54]. Biochemical constraints likely underlie the extensive co-option and repeated evolution in CG biosynthesis, highlighting the critical role of P450 gene turnover in driving diversification [50].

By reconstructing the ancestral state of biochemical plasticity in the tribe Heliconiini, we demonstrated that plasticity was ancestral in the tribe. Yet, some specific clades have undergone specialization either in CG biosynthesis (genus *Eueides* and the clade Doris) or sequestration (Erato clade), and even virtually lost biochemical plasticity (Aoede and Sapho clades). This derived accommodation of plasticity suggests that the dual strategy system used by Heliconiini to acquire their toxins represents a novel example of plasticity-first evolution - in which ancestral plasticity allows colonization of new environments / ecological niches, but the alternative phenotype can be tuned (genetic accommodation) or even become fixed (genetic assimilation/canalization) as lineages specialize in the novel conditions. Although the ability to biosynthesize CGs allowed Heliconiini butterflies to maintain their toxicity regardless of the CG composition of their larval hostplants, widening their diet-breadth, as lineages evolved preference for a specific hostplant selection favoured specialization towards one biochemical strategy over the other. The loss of plasticity in the Sapho clade is associated with the disruption of their *CYP405*, which lacks regions that code for some of the conserved motifs that are crucial for the catalytic activity of a cytochrome P450. Species in this clade tended to have several incomplete copies of this gene and only sequester CGs. We also identified episodic diversifying selection on CYP405 and CYP332 in the Sapho clade. When a plastic phenotype is exposed to selection more often, there is greater opportunity for adaptive refinement [55]. Supporting this, Sapho clade species tend to be monophagous and have preference for a small group of *Passiflor*a in which CG for sequestration is common [56], which could have favoured the loss of biochemical plasticity in this lineage.

To date, examples of plasticity-first evolution in which the associated genetic changes are understood are scarce. For example, snowshoe hares (*Lepus americanus*) in North American often have white colouration during winter and molt to brown in spring[57]. This plasticity is ancestral in the species, allowing the hares to have an optimal camouflage in different seasons. Nevertheless, some populations in warmer areas, such as British Columbia, lost this plasticity and always display a winter-brown phenotype. This loss of plasticity via genetic assimilation/canalization was caused by the introgression of an *Agouti* allele from black-tailed jackrabbits to this population of snowshoe hare [57]. Similarly, side-blotched lizards (*Uta stansburiana*) living on the Pisgah Vulcan are cryptic and have a plastic colouration [58]. The Pisgah phenotype is melanic and inhabits the lava flow, whereas the off-lava phenotype is lighter and cryptic in nearby light-coloured soils. Although both lizard populations can produce the on- and off-lava phenotypes, genetic accommodation has led them to specialize on the phenotype that is most cryptic in their local environment. It has been shown that the Pisgah population can develop darker coloration than the off-lava populations on dark substrate, because they have specific variants of the genes *PREP* and *PRKAR1A* which control melanin formation [58]. These are two clear examples of the importance of ancestral plasticity in morphological traits for the colonization of new environments, as well as the role of refinement in specific lineages towards local specialisation for diversification. Our results on biochemical plasticity in Heliconiini butterflies provide another example of plasticity-first evolution in which the genetic basis for the trait is understood.

Phenotypic plasticity requires a molecular switch mechanism for sensing the environment and controlling the expression of alternative phenotypes[2]. While biochemical plasticity is controlled by the presence of CG for sequestration in the larval hostplant, how butterflies perceive these compounds in their diet and modulate the level of CG biosynthesis in response to that remains unknown. The vast majority of *Heliconius* clades evolved to have at least two complete *CYP405As.* We initially hypothesized that these were redundant orthologues, but our CRISPR experiments suggest that both *CYP405A4* and *CYP405A6* are essential for CG biosynthesis in *H. erato*. We predict that the multiple *CYP405* copies might be necessary to fine-tune this pathway according to diet and modulate biochemical plasticity. In fact, the metabolism of toxins is often composed of enzymes that though very similar, are not redundant and are involved in modulating responses to the environment. For example, CYP83A1 and CYP83B1 in *Arabdopsis thaliana* are very similar and catalyse the same step in the biosynthesis of glucosinolates [59,60]. Despite their similarity, CYP83A1 is more efficient in using aliphatic oximes, whereas CYP83B1 is better for aromatic oximes, and the two orthologues enable *A. thaliana* to precisely modulate their glucosinolate profile in response to stress and developmental changes [59,60]. To detoxify glucosinolates from their Brassicaceae hostplants, *Pieris* caterpillars utilize the enzymes NSP (Nitrile-Specifier Protein) and MA (Major Allergen) that convert these compounds into inert nitriles [61]. Although NSP and MA derived from the same ancestor gene, NSP expression is upregulated by high concentrations of aliphatic glucosinolates, in contrast with MA that is upregulated by aromatics[62]. It will be interesting to determine whether *Heliconius CYP405As* show similar complementary functions.

In summary, we have provided the first functional evidence for the genetic basis of CG biosynthesis in heliconiine butterflies and performed ancestral reconstruction analyses on their biochemical plasticity. Our results bring empirical evidence for an example of plasticity-first evolution, with plasticity ancestral in the tribe and virtually lost in one clade specialized on sequestration. This loss was associated with genes involved in CG biosynthesis losing key functional domains.

## Supporting information

Chemical dataset

## ACKNOWLEDGMENT

We thank the Zoology Department for hosting this project, especially Glennis Julian who maintained the butterfly and plant stocks used in the CRISPR experiment. We also thanks Mika Zagrobelny and René Feyereisen for their amazing work that set the foundation for this project, as well as discussions over the years about the evolution of cyanogenesis in Lepidoptera. We thank the staff at the section for Plant Biochemistry, Department of Plant and Environmental Sciences, University of Copenhagen, for the assistance in the target-metabolomic analyses, especially Mariela Alejandra González Ramírez for the maintenance of the analytical instruments and Mohammed Saddik Motawie for the chemical standards used in these analyses. We thank David Nelson for naming the P450s identified in this project. EC is thankful for the H2020 Marie Skłodowska-Curie Actions (841230) which funded this project. EC, CJ and IW thank the financial support from UKRI – NERC (NE/W005131/1). FC and SHM thank the financial support from UKRI - NERC (NE/N014936/1) and ERC (758508).

## SUPPLEMENTARY METHODS

### Butterfly husbandry

For the CRISPR experiments, fresh eggs were supplied from a insectary population of *Heliconius erato demophoon* keep at the Zoology Department. Butterflies are kept in the green house at 28 °C degrees, 80% Relative humidity and 12h/12h light cycle and fed *ad libitum* with artificial nectar as well as flower of *Lantana camara* and *Psiguria sp*. Plants of *Passiflora biflora* with fresh tips are provided for egg laying and eggs are collected to be raised in cages in a CT room at 28 °C degrees, 80% Relative humidity and 12h/12h light cycle. Larvae are fed *ad libitum* with fresh shoots of *P. biflora* and pupae are transferred to a pupal cage (40 x 40 x 60 cm) for eclosion. Freshly emerged butterflies are transferred to the green house for breeding.

### Classification of CGs

**Table S1.**
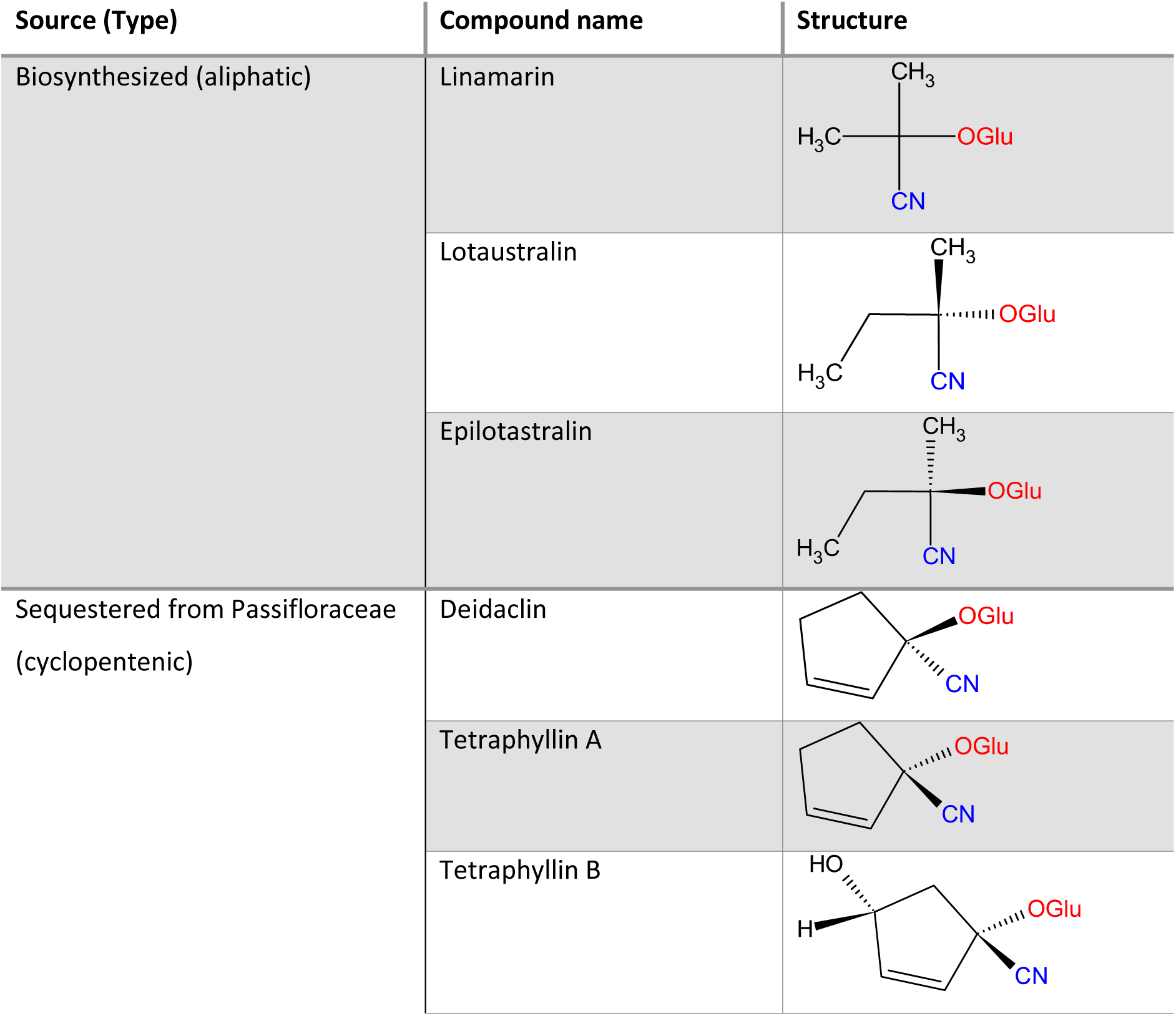
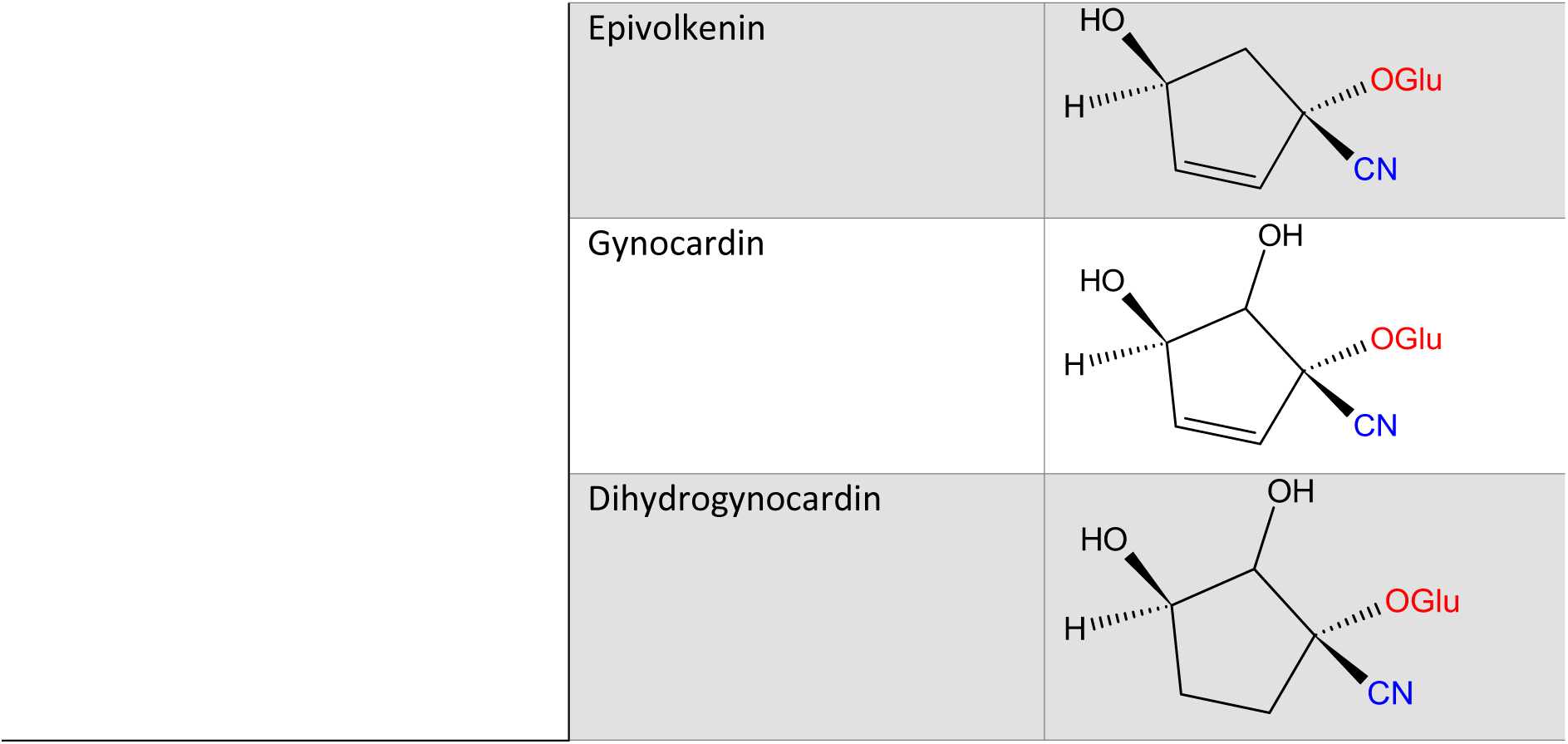
Cyanogenic glucosides (CGs) de novo biosynthesized and sequestered by heliconiini identified in de Castro et al. (2019), Sculfort et al. (2020) and Rueda (2023).

### Statistical Analyses

The larval weight of all injected individuals did not have a normal distribution (Shapiro-Wilko, W = 0.91115, p-value = 1.956e-08). Thus, Kruskal-Wallis was used to investigate if differences in larval weight were significant, followed by Dunn’ Test for pairwise comparison. The CG concentration of larvae that were successfully phenotyped and genotyped also did not have a normal distribution. Kruskal-Wallis and Dunn’ Test were used to evaluate if the differences in CG concentration between control, non-edited (-) and edited larvae (+) were significant.

## SUPPLEMENTARY FIGURES

**Figure S1.**
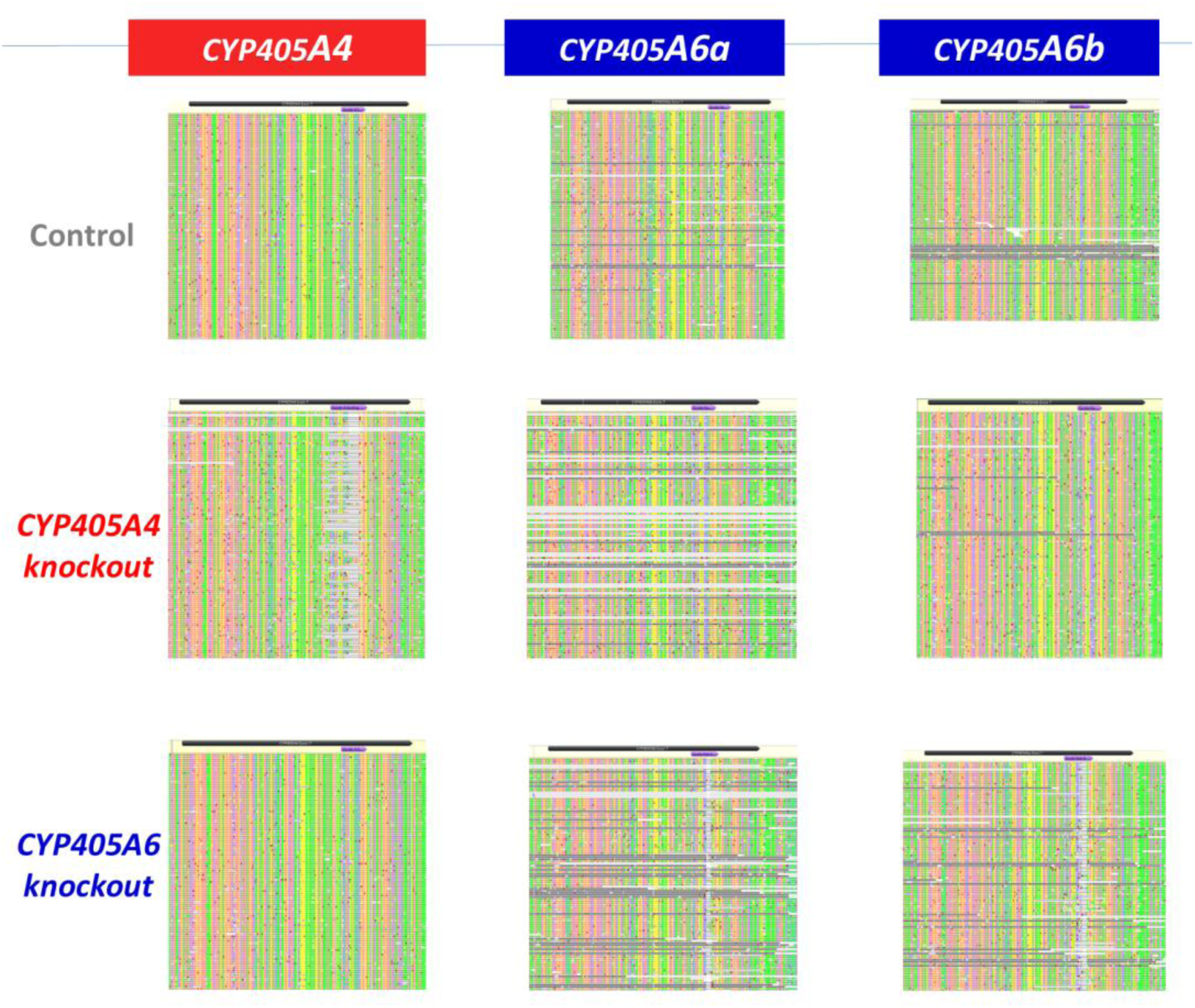
Each row correspond to a single genotyped *H. erato demophoon* individual and each column to a *CYP405* orthologue rectangle. Alignments show Nanopore sequencing results of exon 7 for each orthologues (black box) with the target-CRISPR (purple box). Deletions (grey mark in columns) around the target CRISPR sites demonstrates that guides RNA were specific for the orthologues they were designed for.

**Figure S2.**
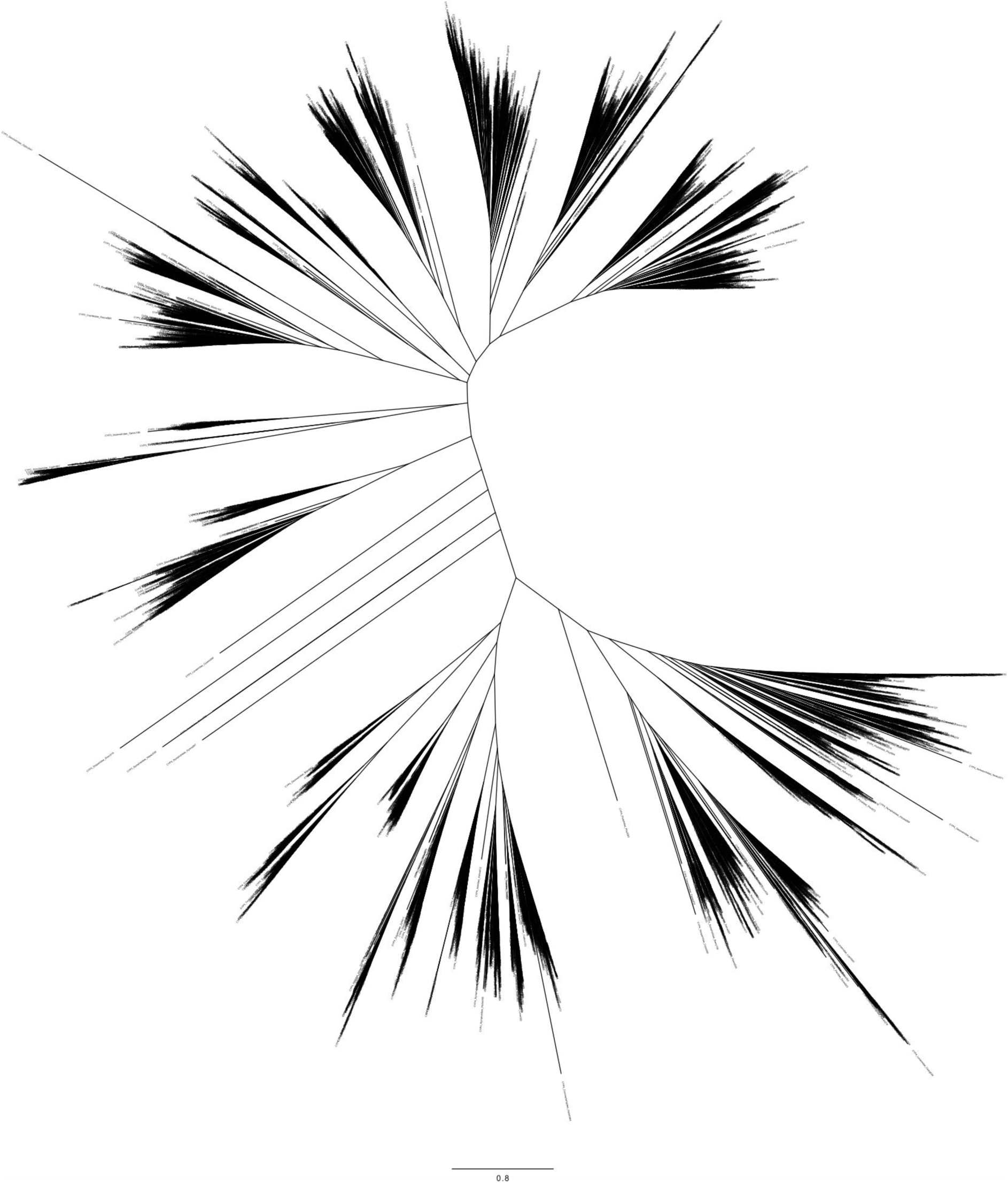
Tree built with the protein sequences of all CYP3 and CYP4 genes in Lepidoptera from insectp450.net database [34] (9,230 sequences) as well as CYP405 and CYP332 in 63 Heliconinae genomes recently published [24] (200 sequences).

**Figure S3.**
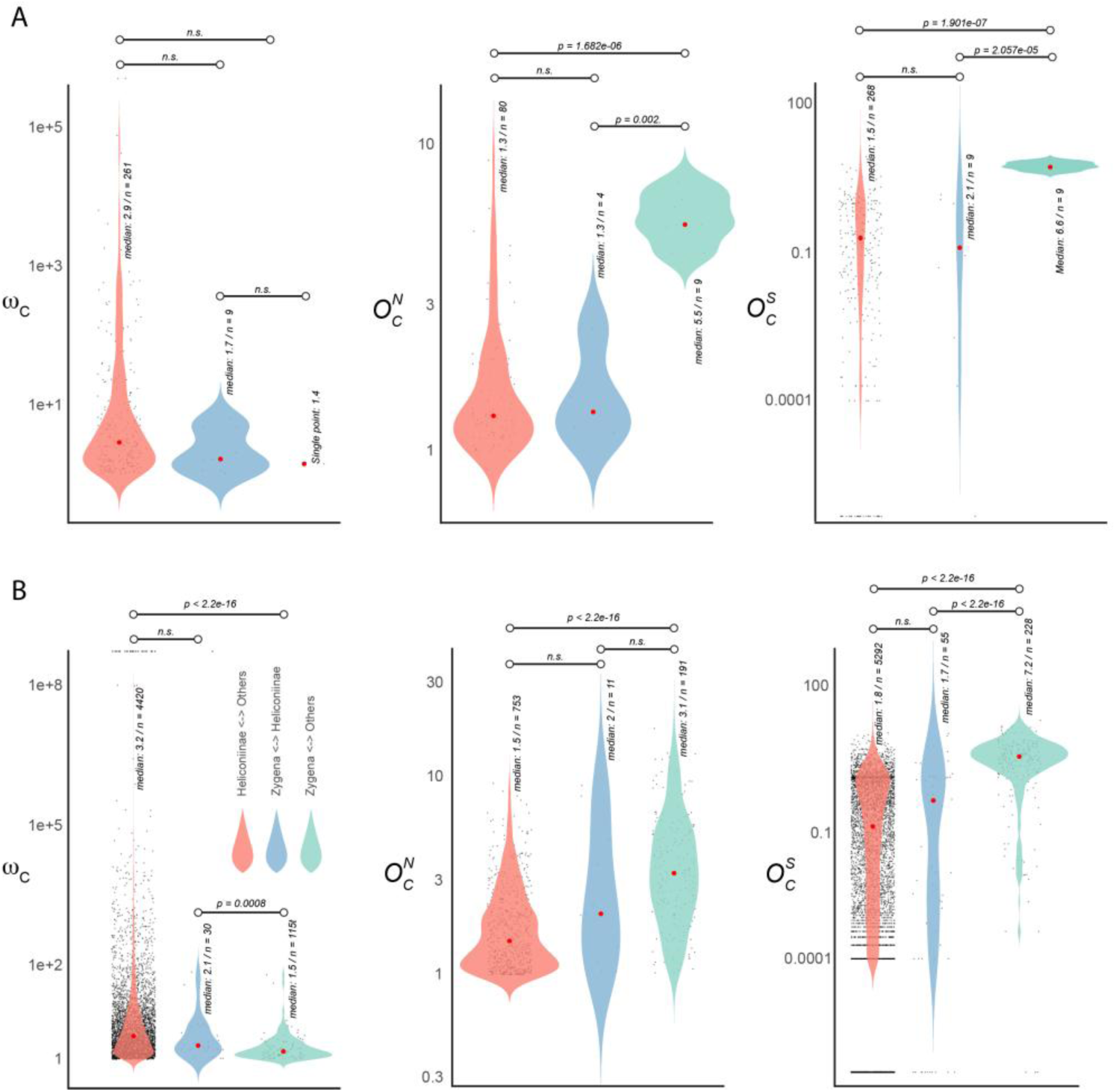
Convergence of positive selection in **A)** *CYP405* and **B)** *CYP332* in Lepidoptera. In the left side the distribution of convergence of positive selection (ω_C_), in the middle and left the rate of observed non-synonymous (O^N^_C_) and synonymous substitutions (O^N^_C_). The violin plot shows three different comparisons: Heliconiinae versus other lepidopteran branches (red); Heliconiinae *versus Zygaena* branches(blue); and *Zygaena versus* all the other lepidoptera branches (green). For *CYP405* only a few branches showed signs of putative convergence, and there was no significant enrichment (one-sided Wilcoxon rank-sum test p-value > 0.05) across the board, with the exception of O^N^_C_ and O^S^_C_, which in both cases are enriched in the comparison of *Zygaena* versus all other lepidopterans, indicating an excess of convergence of non-synonymous sites that is likely stochastic and not due to selection. For CYP332 the pattern was very different: *Zygaena* shows a higher enrichment of ω_C_ in the comparison with Heliconiinae (median 2.1), rather than with all the other Lepidopterans (median 1.5, one-sided Wilcoxon rank-sum test p-value = 0.0008), which also shows a lower enrichment in convergence of positive selection with the comparison of Heliconiinae *versus* all the other lepidopterans (median 3.2, one-sided Wilcoxon rank-sum test p-value < 2.2e-16). Interestingly, there was an excess of non-synonymous substitutions in CYP332 for the comparison between *Zygaena* versus all the other Lepidopterans (Figure 6b). In summary, these analyses suggests that both *CYP405* and *CYP332* were independently co-opted into CG in *Zygaena* moths and Heliconiinae butterflies with little evidence for convergence at the molecular level.

